# Uncovering sex differences in Parkinson’s Disease through metaanalysis of single cell transcriptomic studies

**DOI:** 10.1101/2024.12.20.628852

**Authors:** Fernando Gordillo-González, Irene Soler-Sáez, Cristina Galiana-Roselló, Tim-Simon Burmeister, Borja Gómez-Cabañes, Rubén Grillo-Risco, Marta R. Hidalgo, Beatriz Dolader-Rabinad, Natalia Yanguas-Casas, Ricardo A Ramirez-Hernandez, Franc Casanova-Ferrer, Silvia Salvador-Guerrero, Marina Romero-Ramos, Francisco García-García

## Abstract

Abundant evidence supports the significant impact of biological sex on various aspects of Parkinson Disease (PD), including incidence, progression, symptoms or response to treatment. The incidence and prevalence of the disease is higher in males, while its age of onset is earlier than in females. There are also sex differences in the symptomatology, both motor and non-motor. In female PD, tremor, pain, depression and dysphagia are predominant, whereas in male PD, freezing of gait, camptocormia, cognitive impairment and urinary dysfunction are more common. Likewise, there are sex differences in the pathophysiology of the disease, related to most of the pathological processes of PD, as is the case of the greater activation of microglia in males or the lower oxidative stress in females. All these findings support the idea that different molecular mechanisms may be involved in PD depending on the sex of the patient. Some explanations for these events are related to biological, genetic, hormonal or environmental factors, such as the possible anti-inflammatory and neuroprotective effect of estrogens. However, the underlying molecular mechanisms have not yet been fully described. Our results show sex differences in gene expression, cell-cell communication and pathway activation in all major brain cell types, highlighting the presence of greater neuroinflammation in men and greater neurodegeneration in the SNpc in men, with the latter appearing to be inverted between sexes when observed in the cortical zone. Finally, we have made all the results available in a publicly accessible webtool, in order to allow the exploration of results to other researchers and to broaden the molecular-level understanding of PD sex differences.

## Introduction

Parkinson’s disease (PD) is the most common movement disorder and second most prevalent neurodegenerative disease after Alzheimer’s; affecting over 6 million individuals globally, with both incidence and prevalence continuing to rise in recent years. (Simon et al., 2020). At the neuropathological level, PD is classically characterized by progressive degeneration of dopaminergic neurons of the substantia nigra pars compacta (SNpc) and the intraneuronal accumulation of alpha-synuclein in the so called Lewy bodies, which are found both in central and peripheral nervous system. Although neuronal loss seems to occur in limited areas such as SNpc, dorsal motor nucleus of the vagus nerve, and nucleus basalis of Meynert (<u>10.3389/fneur.2018.00455</u>) it is accepted that the disease involves the neuronal dysfunction in other areas where alpha-synuclein pathology occurs, such as the cortex. While the brainstem is primarily affected in earlier disease stages, the pathology progressively extends to cortical regions in later stages (Braak et al., 2003; Del Tredici & Braak, 2016). This extensive neurodegeneration leads to the onset of both motor (eg. Bradykinesia, rigidity, postural instability) and non-motor (eg. Sleep problems, hyposmia, anxiety, dementia, depression) symptoms of the disease with no treatment capable of decreasing the progression of PD to date (Bloem et al., 2021; Tolosa et al., 2021).

Despite research efforts, the causes behind PD are not yet fully described. However, researchers have been able to identify molecular hallmarks for this disease, including failure of protein clearance pathways, mitochondrial damage, oxidative stress, excitotoxicity and neuroinflammation (Dong-Chen et al., 2023; Maiti et al., 2017; Panicker et al., 2021). Additionally, several risk factors have been implicated in the development of sporadic PD, which accounts for approximately 85% of cases, including advanced age, multiple genetic variants, pesticide exposure, and sex (Belvisi et al., 2020; Cerri et al., 2019; Vázquez-Vélez & Zoghbi, 2021).

Among these factors, sex has received increasing attention due to its multifaceted influence on both clinical presentation and disease trajectory. Evidence suggests that sex differences significantly impact disease incidence, progression, symptomatic presentation and treatment outcomes. Males exhibit a higher prevalence of PD and tend to develop symptoms at an earlier age than females (DuMont et al., 2023; Picillo et al., 2017). Further, sex-related differences have been reported in both motor and non-motor symptoms of the disease. In female individuals with PD, tremor, pain, depression and dysphagia are predominant, whereas in males, freezing of gait, camptocormia, cognitive impairment and urinary dysfunction are more common (Cerri et al., 2019; Meoni et al., 2020). In parallel, sex-differences in the pathophysiological mechanisms have been also suggested, such as a higher activation of microglia in males or the lower oxidative stress in females (Barko et al., 2022; Cerri et al., 2019; Nicoletti et al., 2023). Taken together, these findings support the hypothesis that sex can impact PD pathogenesis through distinct molecular mechanisms. While these differences have been related to distinctive biological, genetic, hormonal or environmental factors -such as the possible anti-inflammatory and neuroprotective effect of estrogens-, the underlying molecular mechanisms have not yet been fully described (Terrin et al., 2023; Vaidya et al., 2021).

A major challenge in PD is the great inter-individual variability symptomatology, disease progression and treatment response. This heterogeneity, coupled with the absence of early biomarkers for diagnosis and monitoring disease progression, complicates clinical management (Bloem et al., 2021). Recent efforts to address this caveat has resulted in several proposals to classify the disease using α-syn related assays, imaging and genetics in different orders and degrees (<u>10.1016/S1474-4422(23)00404-0</u>, <u>10.1016/S1474-4422(23)00405-2</u>), which has ignited a debate in the PD research and clinical community. These classifications are the result of the effort to achieve a biological classification of the disease and a more personalized medicine. In this effort, the study of sex differences would have an important impact as the evidence supporting distinctive molecular and cellular changes among sexes in PD are accumulating. Indeed, in recent years some candidates have been proposed as sex-specific biomarkers (serum urate levels as inverse predictor of PD in male or hypertriglyceridemia as a predictor of cognitive impairment in women, among others), although the data are still far from conclusive and require further validation in larger, stratified cohorts (Ascherio et al., 2009; Luca et al., 2022; Nicoletti et al., 2023). To address this problem, high-throughput technologies, such as single cell RNA-seq (scRNA-seq), allow us to assess these differences in a more global and comprehensive manner. In addition, by integrating the numerous omics studies available in public repositories, the difficulty of inter-patient variability can also be overcome.

Built on the aforementioned, Here we integrated six single-cell transcriptomic studies performed on postmortem brain tissue from individuals with PD, together with two cell-type-specific meta-analyses conducted across distinct brain regions, to improve the resolution and comparability of molecular features across studies and areas. To integrate data, we use the meta-analysis strategy, which increased statistical power and reduced inter-study variability. Our results show sex differences in gene expression, cell-cell communication and pathway activation in all major brain cell types. Notably men exhibited greater neuroinflammation and neurodegeneration in the SNpc. Conversely the pattern of neurodegeneration in the cortical areas appeared reversed in females compared to males suggesting a region-specific and sex-dependent pathological processes. Finally, to enable further analysis and exploration of sex specific molecular mechanisms in PD we developed a publicly accessible web-based tool that allows interactive exploration of all results.

## Materials And Methods

The bioinformatics processing and analysis was carried out using the R programming language (R Core Team, 2021) (version 4.1.2). The packages and versions used are listed in Table S1.

### 1. Study selection

A systematic review was conducted following the Preferred Reporting Items for Systematic Reviews and Meta-Analyses (PRISMA) criteria (Liberati et al., 2009; Page et al., 2021), whose main objectives are standardization and reproducibility. First, we searched public repositories such as GEO (Gene Expression Omnibus) (Edgar et al., 2002), ArrayExpress (Athar et al., 2019), BioStudies (Sarkans et al., 2018), ENCODE (Y. Luo et al., 2020), UCSC Cell Browser (Kent et al., 2002), Human Cell Atlas (Regev et al., 2017), and Single Cell Expression Atlas (Moreno et al., 2022) using the keywords: "Parkinson’s", "Parkinson’s disease" or "PD". To identify suitable datasets, we set the following inclusion criteria: i) organism: *Homo sapiens* and, ii) study type: single-cell or single nuclei RNA sequencing (scRNA-seq and snRNA-seq). Among the selected studies, we discarded those that met any of the established exclusion criteria: i) experimental design not involving individuals with PD versus controls, ii) not focused on PD, iii) lack of information on patient sex, iv) lack of representation of both sexes in all conditions, and v) samples not from post-mortem brain tissue (fetal, organoid, etc.). Finally, the raw count data and associated metadata of the selected studies were downloaded either manually or using the R package GEOquery (Davis & Meltzer, 2007).

### 2. Individual analysis

All selected studies were individually analyzed following three main steps: snRNA-seq data processing, cell type annotation, and differential expression analysis. We performed these analyses based on the guidelines proposed by Amezquita et al., which are based on the use of packages from the Bioconductor repository (Amezquita et al., 2020).

#### a. snRNA-seq data processing

All downloaded data from the different studies (count matrix, cell data, patient metadata, etc.) were stored in a single SingleCellExperiment (SCE) using the SingleCellExperiment R package (Amezquita et al., 2020), or the read10xCounts function of the DropletUtils R package (Lun et al., 2019), depending on the available data format. Afterwards, we performed a homogenization of the nomenclature used for the genes to enable integration between studies.

We then perform quality control to discard droplet counts not derived from a single viable cell. To detect the empty droplets, we used the original paper thresholds for the number of Unique Molecular Identifiers (UMIs) and the number of detected genes. We also discarded those droplets with a percentage of mitochondrial genes higher than 5%. Genes with less than 5 counts were also removed. Finally, we considered as doublets those droplets identified by the scDblFinder R package, as well as those with a library size larger than 4 MADs (Mean Absolute Deviation), since such abundant gene expression is not expected in brain cells. Normalization was performed using the scran (Lun et al., 2016) and scater R packages.

We selected the top 25% of the most variable genes and performed a principal component analysis (PCA) for dimensionality reduction, using scran and scater packages respectively. We next used the PCAtools R package (Blighe & Lun, 2021) to select the number of principal components (PCs) that summarize the largest amount of variance. These selected PCs were used for the T-distributed Stochastic Neighbour Embedding (tSNE) and the Uniform Manifold Approximation and Projection (UMAP) constructed for visualization purposes. Finally, we performed cell clustering using the scran package, applying a combination of the k-means and Shared Nearest Neighbours algorithms. The number of centroids and neighbours were adapted to the number of cells in each study, in order to maintain a similar resolution between them.

#### b. Cell type annotation

Major cell types were annotated combining unsupervised strategies to increase robustness and avoid individual analysis bias. We integrated annotation approaches based on full transcriptomic references and marker gene expression profiles.

For the reference expression profiles-based approach we use the singleR (Aran et al., 2019) and CHETAH (de Kanter et al., 2019) packages, using in both cases as reference dataset a subset of the Allen Brain Map atlas (Bakken et al., 2021), adapted to the recommendations of each software. For the marker gene approach, we used the SCINA R package (Z. Zhang, 2019), using the reference markers present in the CellMarker database (X. Zhang et al., 2019), and BRETIGEA R package (McKenzie et al., 2021), using the package’s own marker genes. Subsequently, to obtain the consensus annotation, we assigned each cell as the most frequent annotation among all the methods, being 2 the minimum number of algorithms that must coincide in the same annotation. Those cells with indeterminate annotation or not annotated as the majority cell type of the cluster were discarded to avoid possible biases in subsequent analyses.

We annotated astrocytes, neurons, microglia and oligodendrocytes in the different studies, as these are the 4 major cell types in brain tissue. Clusters that were not correctly annotated were discarded for the visualization of results and they were not used in the downstream analyses.

Finally, the expression of marker genes tested in the original studies were evaluated using the scater’s plotExpression function to check that the annotations were performed correctly.

#### c. Differential expression analysis

Differential expression analyses (DEA) were performed using the MAST R package (McDavid et al., 2021). First we built a hurdle model with the zlm function, giving as the main variable the combination of condition and sex. We also used the sample that each nucleus came from to reduce intra-patient bias. With this model, we used functions lrTest to obtain the p-values and getLogFC to obtain the logarithm of fold change (logFC) and the standard deviation (SD) of the genes. To obtain the sex-based differentially expressed genes (DEGs), we applied the following contrast:

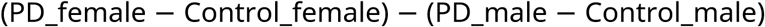

This contrast reveals the sex differential impact of disease, identifying those genes whose expression contributes to the pathophysiology of the PD in a manner that differs depending on the sex of the patient, taking into account the basal differences at the physiological level that may exist between healthy males and females.

In this context, a positive logFC in this contrast could mean that, during PD progression, the gene: i) is upregulated in females and downregulated in males; ii) is upregulated in both sexes, but the increase in expression is higher in females than in males; iii) is downregulated in both sexes and the decrease in expression is lower in females. For simplicity, we refer to these genes as increased in females. Otherwise, the opposite is interpreted for genes associated with a negative logFC, which we designate as increased in males.

#### d. Signaling pathway activation analysis

The hipathia R package (Hidalgo et al., 2016) was used to compute the activation values of the KEGG signaling pathways from the normalized gene expression matrices of the individual single cell datasets. This software divides each pathway into distinct sub-pathways associated with a particular end node or effector of the pathway, obtaining an activation value for each one. Thus, with this step we obtain the activation levels of each sub-pathway in each cell of the dataset. Subsequently, we perform a differential activation analysis (DAA) using MAST and the sex-differential double contrast as described in the previous section. We considered differentially activated sub-pathways (DA-subPs) those with a FDR lower than 0.05.

### 3. Meta-analysis

DEA and DAA results were integrated into 2 meta-analyses, one including datasets of the cortical region and other of the nigrostriatal pathway. The meta-analyses were performed using the R package metafor (Viechtbauer, 2010), using the DerSimonian and Laird random effects model, which accounts for the heterogeneity of each individual study, giving greater weight to those studies with lower dispersion. These analyses calculate for each gene its p-value, logFC and SD among other statistics. The p-value was corrected using the Benjamini & Hochberg (BH) algorithm, and those genes with a p-adjusted value < 0.05 and a logFC > 0.5 were considered significant DEGs. In the case of pathways, they were considered DA-subPs when the p-adjusted p-value was lower than 0.05. Finally, we applied diagnostic tests on the obtained results to avoid possible bias effects, including graphical representation of DEGs with funnel and forest plots, and influence and sensitivity analyses (leave-one-out cross-validation).

### 4. Functional enrichment

We performed a biological characterization for each cell type in both sexes from its increased DEGs. The weight01 algorithm (Alexa et al., 2006) from the topGO R package (Alexa & Rahnenfuhrer, 2023) was used for the Biological Process (BP) GO ontology with a minimum node size of 15 genes.

The significant GO terms obtained (p-value < 0.05) were further processed with the R package simplifyEnrichment, to obtain clusters of similar GO terms, which summarize and facilitate interpretation. For better visualization, these clusters are represented in a similarity heatmap with their associated word clouds. Also, to facilitate the visualization and understanding of the data, we selected the ancestral GO term that best covered the information of each cluster. To select it, we took into account the frequency of appearance of the ancestor within the cluster and its probability of appearance in the GO database, an index obtained using the GOSemSim package.

Protein-protein interaction (PPI) networks were created with the STRINGdb R package, using a threshold score of 400. These networks were then analyzed and modified through the STRING web tool (Snel et al., 2000; Szklarczyk et al., 2020).

### 5. Cell-Cell Communication analysis

Cellular communication between the different cell types was analyzed using the iTalk R package (Y. Wang et al., 2019). We used Omnipath intercell resource, from the OmnipathR R package (Türei et al., 2021), as the database for ligand-receptor interactions, since it integrates most of the available resources of ligand-receptor databases, and also has a curated specific category for intercellular communication analysis. We also used this resource to tag each sender and receiver protein with its feature and functional annotations.

For the analysis, we used DEGs obtained from the meta-analysis as input, assuming as sex-specific interaction gains those whose ligand and receptor are both significant in the meta-analysis for that sex. We used the product of the logFC from the sender and receiver as a measure of intensity in the gain of interaction. We also used the logFC for the representation of the results in circus plots.

### 6. Web platform

We developed a user-friendly online resource to easily explore all the results obtained from the different analyses conducted in this work. The web app (https://bioinfo.cipf.es/cbl-atlas-pd/) was developed under the structure of the R shiny package (Chang W, 2024) and is compartmentalized into the different steps performed in the analysis.

## Results

### Study selection and individual analysis

The systematic review identified 84 unduplicated studies (Figure 1A). Of these, seven studies remained after the established inclusion and exclusion criteria were applied (further details in material and methods section). After quality control analyses, one of these seven (GSE184950 (Q. Wang et al., 2024)) was discarded due to the high expression of mitochondrial genes in the majority of the cells, which is indicative of poor cell viability. Therefore, 6 studies (Table 1) with a total of 110 samples, were included in the final analysis. Sample distribution by sex and condition is reported in Figure 1B. Of these six studies, four were from the substantia nigra (Kamath et al., 2021; A. J. Lee et al., 2022; Smajić et al., 2021; Martirosyan et al., 2025) and two from anterior cortical areas (1 from the anterior cingulate cortex and 1 from the prefrontal cortex) (Feleke et al., 2021; Zhu et al., 2022).

**Figure 1.**
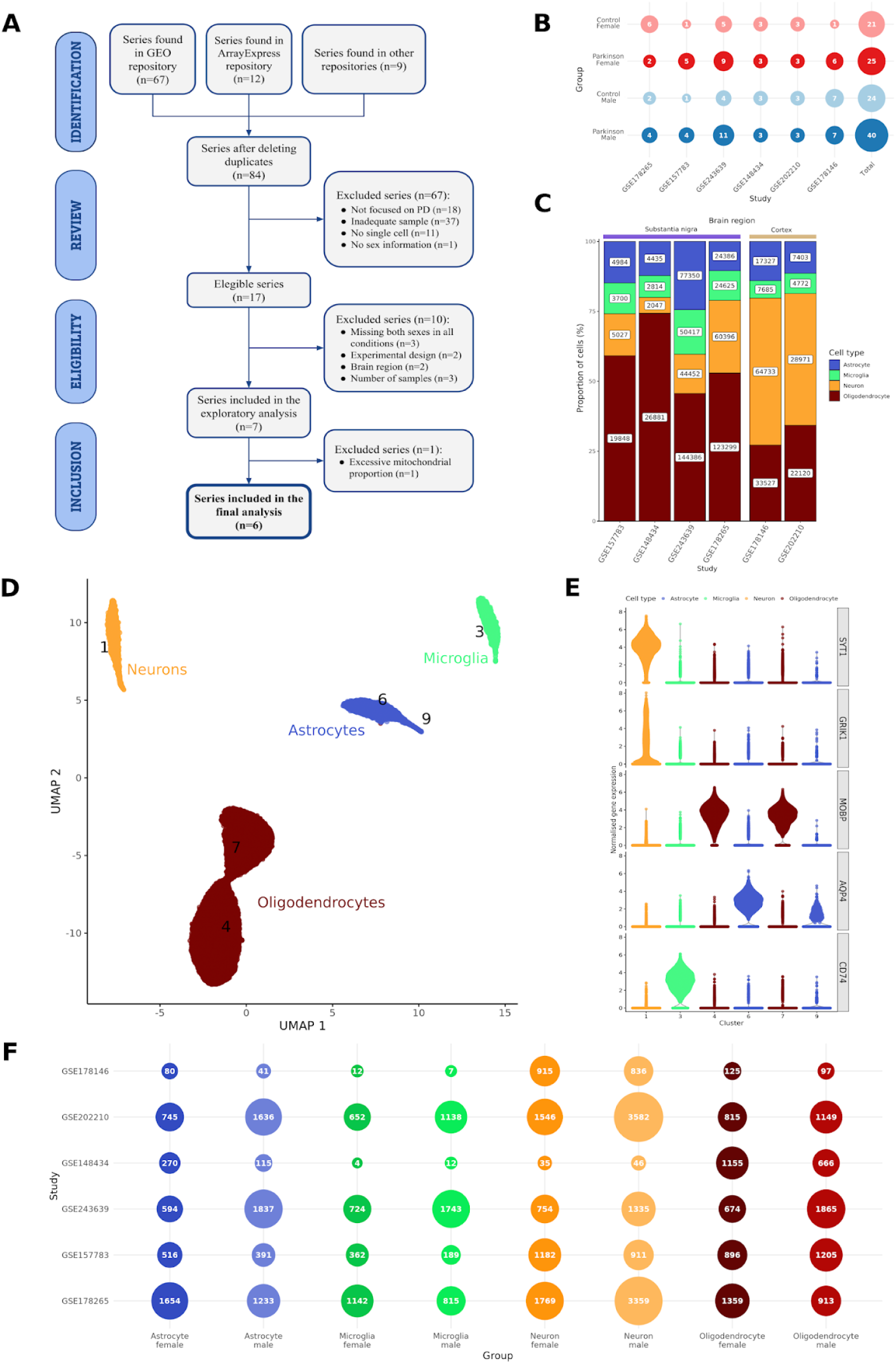
Systematic review and individual analyses results overview. A-PRISMA diagram with the description of the different steps performed for the systematic review. B-Number of individuals from each selected study, grouped by sex-disease condition. C-Barplot showing the proportion and total number of each cell type identified by study. D-UMAP embedding of GSE157783 study, colored by cell type. Numbers correspond to clusters. E-Violin plot representation of the expression of cell type markers for each of the clusters of the GSE157783 study, colored by cell type. F-Number of significant DEGs in each individual differential expression analysis, classified by cell type and sex in which the gene is increased. *DEGs: differentially expressed genes, PD: Parkinson’s disease,* PRISMA*: Preferred Reporting Items for Systematic Reviews and Meta-Analyses, UMAP: Uniform Manifold Approximation and Projection*.

**Table 1.** Description of the selected datasets. * Control female: case female: control male: case male. SNpc: Substantia nigra pars compacta; ACC: Anterior cingulate cortex; PC:

| Dataset | Sequencing method | Sequencing platform | Brain region | Tissue | Patients* | Cell number | Gene number | Genome aligned | Ref |
| --- | --- | --- | --- | --- | --- | --- | --- | --- | --- |
| GSE178146 | 10x Genomics | Illumina HiSeq 4000 | Cortex | ACC | 1 : 7 : 6 : 7 | 156.577 | 25.907 | GRCh38 |  |
| GSE202210 | 10x Genomics | Illumina NovaSeq 6000 | Cortex | PC | 3 : 3 : 3 : 3 | 232.155 | 27.253 | GRCh38 |  |
| GSE178265 | 10x Genomics | Illumina NovaSeq 6000 | SN | SNpc | 6 : 2 : 2 : 4 | 253.029 | 24.020 | GRCh37 |  |
| GSE157783 | 10x Genomics | Illumina NovaSeq 6000 | SN | Midbrain | 1 : 1 : 5 : 4 | 39.110 | 23.383 | GRCh38 |  |
| GSE243639 | 10x Genomics | Illumina NovaSeq 6000 | SN | SNpc | 5 : 4 : 9 : 11 | 316.605 | 27.335 | GRCh38 |  |
| GSE148434 | 10x Genomics | Illumina NovaSeq 6000 | SN | SNpc | 3 : 3 : 3 : 3 | 41.455 | 20.779 | GRCh37 |  |

Prefrontal cortex.

After performing the individual dataset processing, we identified neurons, astrocytes, microglia and oligodendrocytes in each study. The distribution of cell types per study is illustrated in Figure 1C. GSE157783 study cell types were represented in a UMAP embedding (Figure 1D) and the expression of marker genes in each cluster was verified (Figure 1E). Similar results were obtained in the other datasets (Figure S1). Finally, differential expression analyses identified DEGs in both sexes for all cell types in all studies. Although the number of DEGs varied greatly between studies (i.e. microglia in GSE148434 and GSE178146 studies). This disparity highlights the importance of statistically integrating the results obtained beyond those that are significant in each study, as applied in the present analysis (Figure 1F).

### Gene expression meta-analysis

We performed two cell-type-specific meta-analyses of gene expression, one from the four studies of the substantia nigra (SN) and another from those two studies of cortical (CT) regions. We categorized these DEGs as increased in females or males depending on whether the logFC obtained from the double contrast was positive or negative, respectively (Figure 2A). In SN, A total of 357 DEGs were found across all cell types (65 in astrocytes, 84 in microglia, 127 in neurons and 81 in oligodendrocytes), distributed between the two sexes. The meta-analysis in cortical regions revealed 1219 DEGs (127 DEGs in astrocytes, 73 in microglia, 920 in neurons and 99 in oligodendrocytes) (Figure 2C). In both meta-analyses, neurons presented the highest number of DEGs, which corroborates the central role of neurons in PD but also suggest a significant impact of sex in the disease on this cell type. To further investigate this, we represented these results in a volcano plot for each brain region (Figure 2B for SN and Figure 2D for CT), labeling those with greater significance and variation between sexes. See Table S2 for all DEGs obtained in each cell types for both meta-analysis and the online resource (https://bioinfo.cipf.es/cbl-atlas-pd/) to explore the results, and found also sex-stratified analyses (DEA in each sex separately).

**Figure 2.**
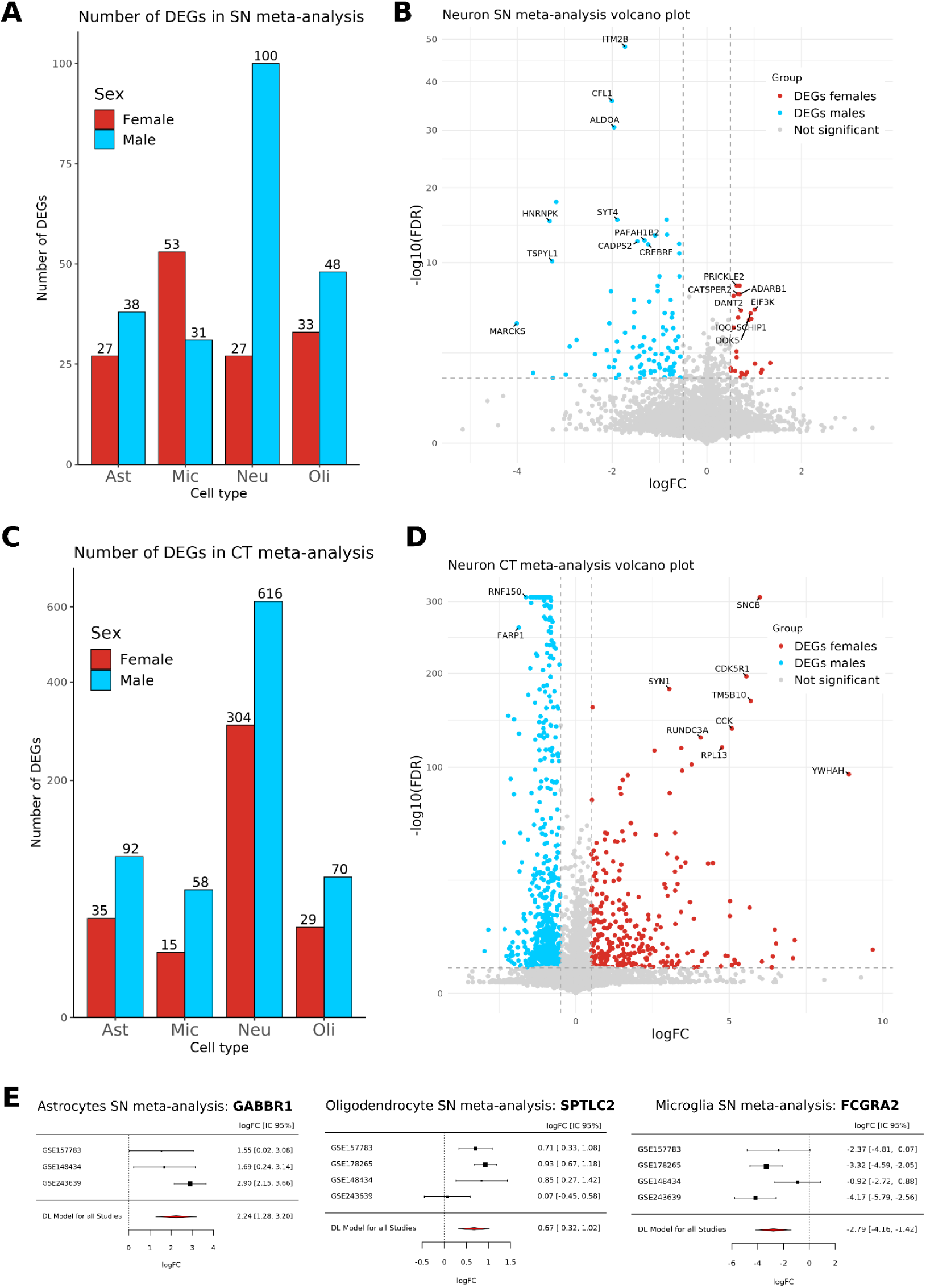
Meta-analyses results of nigrostriatal pathway region. **A,C**-Number of DEGs obtained in the SN (**A**) and CT (**C**) meta-analyses for each cell type, divided by sex. **B,D-** Volcano plot representation of the results obtained in the SN (**B**) and CT (**D**) meta-analysis of neurons. The x-axis represents the logFC of the comparative and the y-axis the -log10 of the adjusted p-value. Those identified as DEGs (FDR < 0.05 & abs(logFC) > 0.5) are colored by sex. **E-** Forest plot of GRIA2 (left), PIBF1 (middle) and VPS35 (right) genes in SN meta-analysis for different cell-types. The forest plots show the logFC and confidence intervals for each individual study and the DL model for all studies (lower red diamond). Ast: Astrocytes, Mic: Microglia, Neu: Neurons, Oli: Oligodendrocytes, SN: Nigrostriatal, C: Cortical, DEGs: differentially expressed genes, FDR: false discovery rate, IC: confidence interval, DL: DerSimonian and Laird.

Interestingly, the genes with highest p-value and logFC included disease relevant genes in both brain regions. In SN, genes related with synaptic release were found, such as SYT4 and CADPS2, whose variants or expression impairments have been related to PD pathology (Smajić et al., 2021, PMID: <u>28647363</u>, PMID: <u>35232250</u>). But also, other genes that might indirectly affect proteins previously highlighted in brain diseases such as ADARB1 -RNA editing enzyme associated with glutamate receptor excitotoxic balance-, and PRICKLE2 - involved in axonal maintenance. Regarding CT, notable genes include SNCB, encoding another protein of the synuclein family that has been shown increased in synucleinopathies and associated with dementia (PMID:<u>35122299</u>), and other synapse-related genes such as FARP1, a postsynaptic signaling and scaffolding protein (PMID: <u>23209303</u>), and SYN1, a presynaptic phosphoprotein that regulates neurotransmitter release (PMID: <u>30065071</u>).

Additionally, we individually explore the forest plots of some potentially PD-relevant DEGs in the SN meta-analyses so as to verify that there were no studies that significantly influenced the overall result of the meta-analysis (Figure 2E). We analyzed the results of GABBR1 in astrocytes, a GABA-B receptor modulating astrocytic Ca²⁺ and gliotransmission ((TIM Ref) PMID: <u>28012274</u>); SPTLC2 in oligodendrocytes, a subunit of serine-palmitoyltransferase (PMID: <u>32041816</u>), which is rate-limiting for sphingolipid/myelin lipid synthesis; and TGFBR2, related with the maintenance of homeostatic microglial phenotype (PMID: <u>40205047</u>).

### Biological characterization of gene meta-analysis

The overrepresentation analysis revealed significantly enriched GO biological process terms in all DEGs sets from the meta-analysis of both brain regions (see number of GO in Figures 3A (SN) and S2A (CT)). The number of GO terms enriched in all cell types in SN are higher in males than in females (Figure 3A). However, in the CT region, while enriched GO terms in neurons are also higher in males, in females more GO terms were found in glia cells and especially for microglia (Figure S2A), despite the starting DEG sets being smaller (Figure 2C). Thus, our analysis suggests a differential affection of cortex or nigrostriatal areas according to biological sex, with cortex being more affected in females and nigrostriatal areas in males.

**Figure 3.**
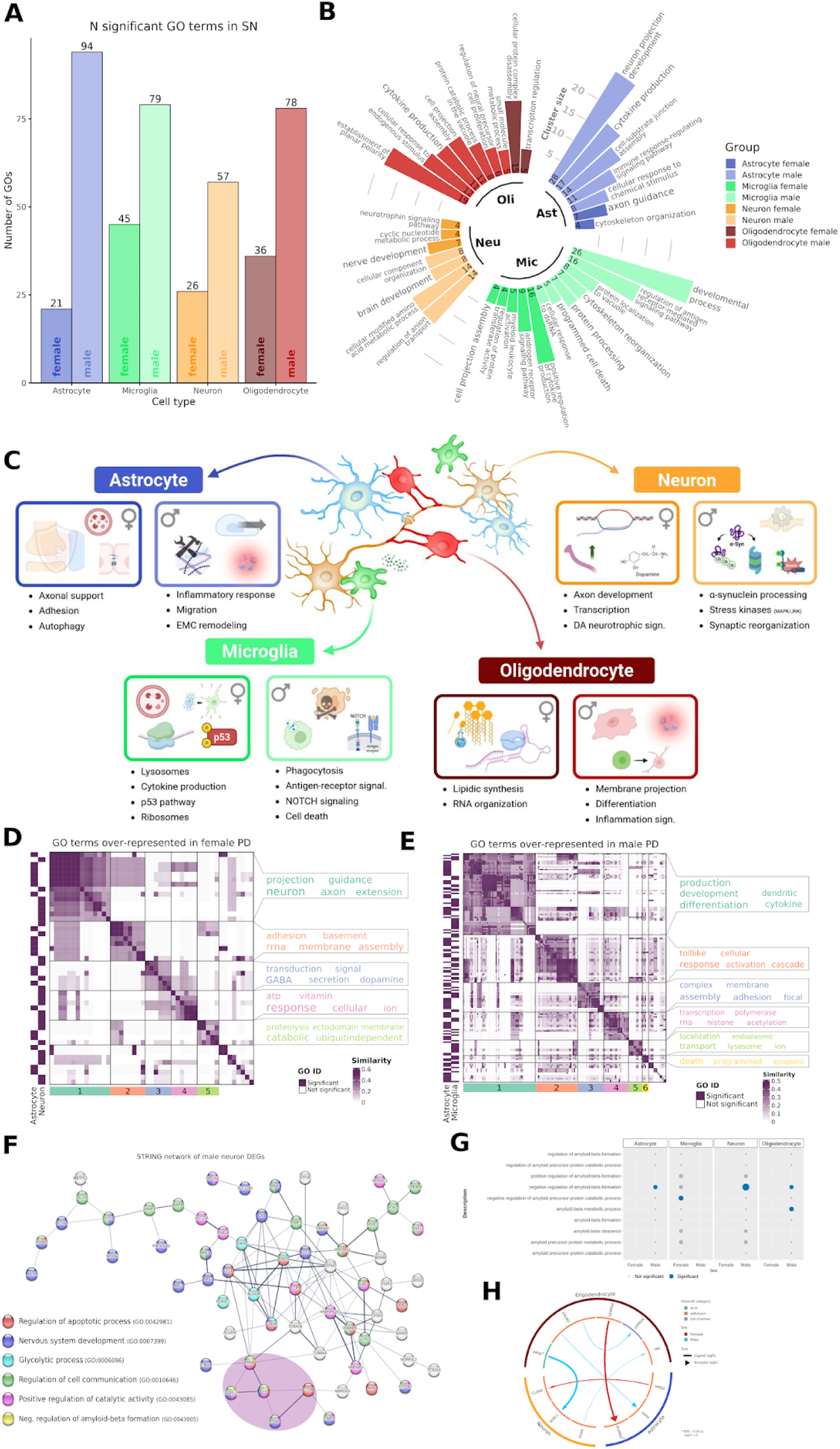
Biological characterization of the results obtained in the meta-analysis of the nigrostriatal pathway. **A-** Number of significant GO terms for each of the cell types separated by sex. **B-** Circular barplot representing the size of the GO term clusters obtained for each of the cell types and sexes. Above the bars, the name of the GO term that best summarizes each of the identified clusters. **C-** Graphical overview of the biological results obtained in each of the cell types for each sex. **D,E-** Heatmap representation of the similarity matrix obtained from GO biological processes enriched in neurons and astrocytes in females (D) and microglia and astrocytes in males (E). Bottom divisions delimit terms grouped in clusters, and on the right side, the word cloud with the most representative words of the cluster. The left margin represents from which cell type each of the GO terms comes from (color indicating presence and blank absence). **F-** Dotplot of GO biological processes related with amyloid-beta processing. **G-** PPI network of the males DEGs obtained in the meta-analysis of neurons. The network was obtained with mean interaction confidence (0.400). Each node is colored by its associated enriched GO-BP. **H-** Circos plot with the cell-cell interactions classified by sender and receiver cell type and by function of each gene in the intercellular interaction. The thicknesses of the body and tips of the arrows correspond to the logFC magnitude of the ligand and receptor, respectively. PPI: Protein-protein interaction, GO: Gene Ontology, BP: Biological Processes, NT: Neurotransmitter, Ast: Astrocytes, Mic: Microglia, Neu: Neurons, Oli: Oligodendrocytes, SN: Nigrostriatal pathway, DEGs: differentially expressed genes, logFC: logarithm of fold change.

For interpretability purposes, these enriched GO terms sets in each cell and sex were grouped based on their semantic similarity, obtaining clusters of terms related to similar functions. Each of these clusters was finally associated with its most representative GO term, based on the frequency of appearance of the ancestor within the cluster and the “rarity” of the GO term (see Methods Section 4), although some of these terms were modified based on their biological relevance in PD. These analyses for all cell types and sexes are shown in Figure 3B (for SN) and S2B (for CT). Significant GO terms categorized by cluster can be found in Tables S3 (SN) and S4 (CT). In addition, the web resource provides more information on this analysis and allows building customized plots on the biological processes of choice (https://bioinfo.cipf.es/cbl-atlas-pd/).

A graphical summary of the main results in the cluster analysis in each cell type in SN is shown in Figure 3C. Differential expression analysis revealed marked sex-specific biological signatures across cell types. In female neurons, DEGs were mainly associated with neurotrophic dopaminergic-related processes and axonal maintenance, including neuron cell–cell adhesion (GO:0007158) and dopaminergic neuron differentiation (GO:0071542). Within these terms, genes such as ASTN2, a synaptic adhesion molecule linked to neuronal connectivity, and PRICKLE2, involved in axonal maintenance, were identified, together with LMX1B, a direct dopaminergic lineage factor. In this context, astrocytes in females showed enrichment for processes related to axonal support and neurotransmission, such as axon extension involved in axon guidance (GO:0048846) and the GABA signaling pathway (GO:0007214). Relevant genes included GABBR1, the GABA-B receptor gene reported to enhance astrocytic morphological complexity and connectivity with neurons, and TECPR2, a gene involved in autophagy, for which variants have been associated with neurodegeneration. To further explore this possible female-specific synergy between astrocytes and neurons, we performed a semantic clustering (see methods 4) of the functional terms overrepresented in these cell types (Figure 3D). The clusters obtained from this analysis corroborate the processes identified individually, highlighting the largest cluster (green), related to the development of neuronal projections, and cluster 3 (blue), with a word cloud related to GABA and dopamine signaling.

Microglia also exhibited clear sex-specific differences. In both sexes, microglial DEGs were enriched in inflammatory-related processes; however, in females these were more strongly associated with cytokine and chemokine production, including positive regulation of chemokine production (GO:0032722). Genes contributing to this enrichment included SYK, a central kinase downstream of Fc/TREM receptors, and HMGB1, an alarmin known to promote cytokine production. In contrast, male microglia showed enrichment for processes related to antigen presentation and phagocytosis, such as regulation of T cell receptor signaling pathway (GO:0050856), with genes including FCGR2A, an Fcγ receptor that drives phagocytosis and pro-inflammatory signaling, and LAPTM5, a lysosomal adaptor linking endocytosis to inflammatory signaling. Notably, closer inspection of the female microglial DEG set revealed the presence of genes previously reported as protective for microglial homeostatic function in disease contexts, including TGFBR2 and RGS10 (REFs), suggesting a more homeostatic function compared to men.

Beyond microglia, male glial populations showed a broader enrichment of inflammatory-related processes. Astrocytes were enriched for regulation of macrophage cytokine production (GO:0010935), while oligodendrocytes showed enrichment for myeloid leukocyte cytokine production (GO:0061082). These processes included genes such as PLCG2, involved in immune and lipid signaling, HSPB1, previously reported as upregulated in glial stress states (REF), and BCL6, a transcriptional repressor described to modulate inflammatory gene networks (REF). We also performed clustering of terms between male microglia and astrocytes (Fig. 3E), as their crosstalk plays an essential role in the neuroinflammation process. The largest clusters (green and orange) obtained show a direct relationship with immune processes such as cytokine production (green) or immune receptor-mediated response (orange). Also noteworthy are cluster 5 (light green), mainly from microglia and related to lysosomes, and cluster 6 (yellow), related to apoptosis.

In addition, protein–protein interaction (PPI) networks were constructed for each DEG set, revealing in most cases significantly interconnected networks. The PPI network of male neurons in SN (Figure 3F) was the largest, consistent with this being both the most extensive DEG set and the central cell type involved in PD. Most proteins formed a large functional module enriched in the nervous system–related processes and cellular communication, including SYT4, SLC12A5, and NRXN2, among others. A highly interconnected metabolic module related to glycolysis was also identified, with genes like ENO1 and ALDOA, suggesting a metabolic shift in male neurons. Of particular interest, we also identified a module related to protein aggregation (purple shade), identified by the significant enrichment of the amyloid beta formation process (GO:0043005) and expanded through literature curation. This module included PRNP, SORL1, CLU, CTSB, and PSAP, all of which have been previously linked to α-synuclein processing, a key mechanism underlying Lewy body formation and PD pathology (REFs).

Finally, further exploration of protein aggregation–related processes showed a sex- and cell-type–specific distribution. Processes associated with amyloid beta aggregation were exclusively detected in female microglia, whereas in males they were enriched across multiple cell types, including neurons, astrocytes, and oligodendrocytes (Figure 3G). This differential distribution highlights distinct cellular patterns associated with protein aggregate– related pathways between sexes. This may suggest that female microglia are more efficient in managing protein aggregates, potentially reducing the burden on other cell types. In contrast, the accumulation of these aggregates in males may lead to a more reactive and stressed cellular phenotype, as other cell types compensate for the inability of microglia to efficiently handle these aggregates. This could contribute to the more pronounced neuroinflammatory state seen in males, further exacerbating PD pathology.

Finally, the cell communication analysis revealed that sex-differential direct interactions were between oligodendrocytes, neurons and astrocytes, being most of them related to adhesion processes. Among the interactions with the highest logFC in ligand and receptor, we highlight APOE-SORL1 in males between oligodendrocytes and neurons, or SEMA5A-PLXNA2 in females between oligodendrocytes and astrocytes (Figure 3H).

The metanalysis in the CT region listed more biological processes. In female neurons, we found GO terms associated with synaptic plasticity, synaptic vesicles, axonogenesis, microtubule reorganization, translation, with the mitochondrial respiratory chain, as well as processes related to cell death. We explore the PPI network for the DEG in neurons, and we found again processes related to ribosomes and to negative regulation of gene expression, neuron system development and a cluster identified as PD related, where mostly genes related to mitochondria the respiratory chain are found (Figure S2C). Astrocytes show significant genes in GO terms related to BBB and with cell adhesion, chemotaxis and cell shape with VEGFA (PMID: 34713920) and PTN (PMID: 35246557) included as significant. Both Astrocytes and oligos showed both regulation to response to stress (oxidative stress for astrocytes) and myelinating processes were carried out by oligos (central nervous system myelination (GO:0022010)).

Regarding males, there are patterns of neuronal survival and synapse strengthening, along with adhesion processes, observed both in individual enrichments (i.e. neurotransmitter reuptake (GO:0098810) and axon guidance (GO:0007411) in astrocytes, neuron projection morphogenesis (GO:0048812) and glutamatergic synaptic transmission (GO:0035249) in neurons, etc.) and in neuron-astrocyte GO term clustering (Figure S2D). Regarding male microglia, we observed processes related to phagocytosis and chemotaxis (positive regulation of phagocytosis (GO:0050766)), highlighting the increase of the CX3CR1 receptor.

Lastly, in the cell-cell communication analysis (Figure S2E), we notice a more important involvement of astrocytes compared with the SN region. Among the interactions with the highest rate of change in females are LIFR-IL6ST between astrocytes and microglia or NRG2-ERBB4 between neurons. Meanwhile, in males, we highlight APLP1-PRNP interaction between neurons and oligodendrocytes, CTTN-ACTB between neurons and microglia or CALM3-CAMK2G between neurons.

### Pathway activation meta-analysis

We further analyzed the differential sex-specific activation patterns of the KEGG signaling pathway, which divides each pathway into distinct sub-pathways associated with a particular effector protein, obtaining an activation value for each one. Using the same double contrast described for the gene meta-analysis, the activation values were compared thus obtaining sex-dependent differentially activated sub-pathways (DAsubPs). With this, we identified sex-dependent activation patterns in all cell types. For synthesis purposes, all the DA-subPs were aggregated by pathways which were categorized based on its associated biological function (Figure 4A, Figure S3A).

**Figure 4.**
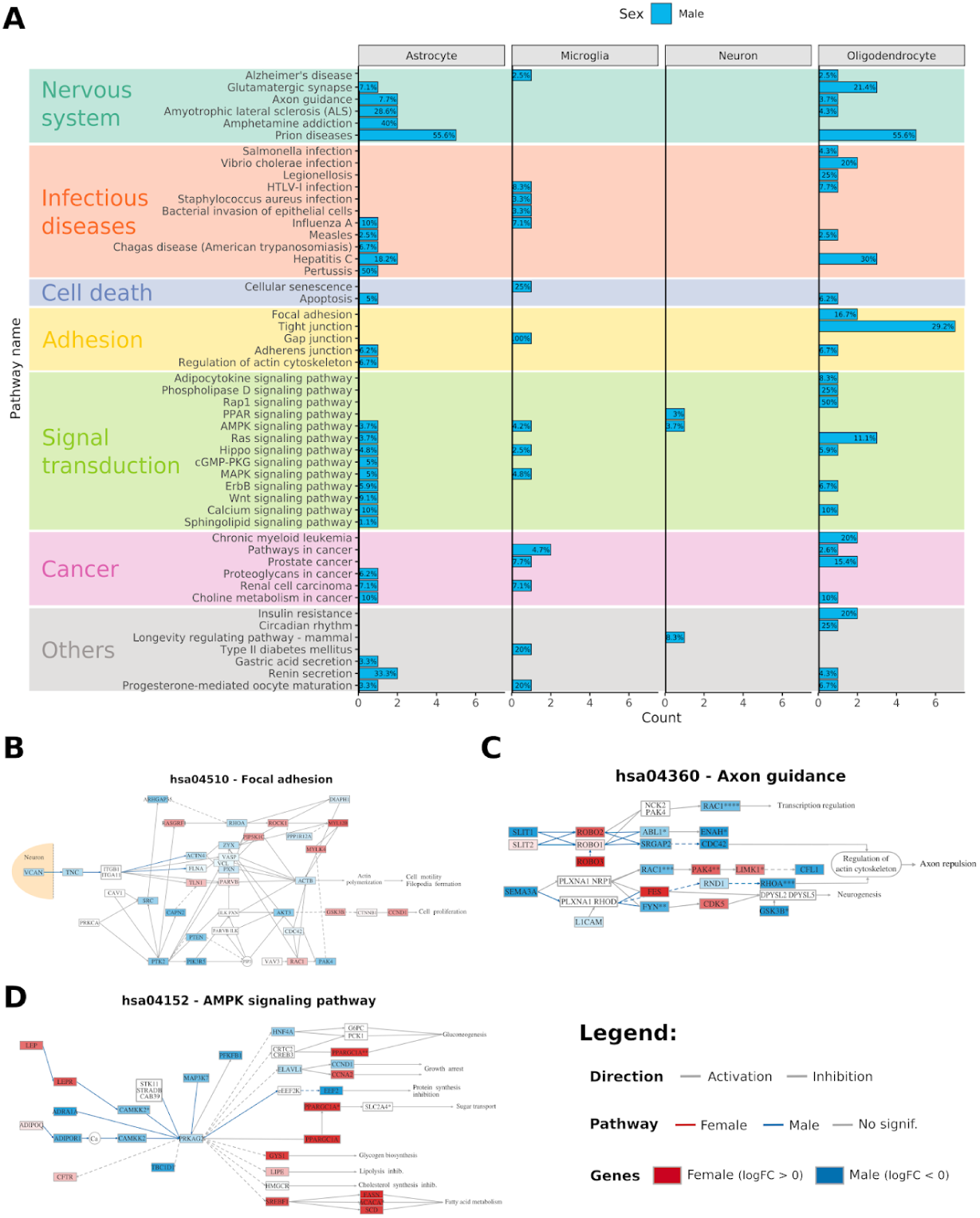
Results of pathway meta-analysis of SN. A- Differentially activated signaling pathways in the meta-analysis of each cell type, coloured by sex. The x-axis indicates the number of significant subpathways and the label inside the bar indicates the percentage over the total number of subpathways in the corresponding pathway. B-D - Graphs representing differentially activated signaling pathways in oligodendrocytes (B), astrocytes (C) and neurons (D). Nodes represent genes and edges and pathways. Coloured edges correspond to significant differentially activated subpathways. The color intensity of the genes is proportional to their FDR in the gene meta-analysis (the lower the FDR, the higher the intensity). To improve visualization, we discarded non-significant subpathways that are not involved in the significant ones.

Regarding the SN, all significant pathways are differentially activated in males, consistently with the higher number of results obtained in this sex on all the analyses (Figure 2A, Figure 3A), probably indicating a higher overall impact of PD in these patients. It is also noteworthy that, in this case, neurons are the cell type with the least significant results, unlike in the previous analysis. This may be because the sex differences observed at the gene level occur in important but isolated genes from various signaling pathways, which may be penalized when detecting differential activation of a complete pathway using this approach. In biological terms, the presence of terms related to nervous system processes is noteworthy, particularly the Prion diseases pathway in astrocytes and oligodendrocytes. Likewise, the presence of pathways related to infectious diseases in all neuroglia cells may be consistent with the reactive phenotype discussed in the previous section. It is also notable the increased presence of adhesion pathways mainly in oligodendrocytes.

To obtain a complete overview of each process and combine these results with the gene meta-analyses, we plotted the more biologically relevant significant subpathways together with the logFCs from the previous gene meta-analysis. It is notable that the focal adhesion pathway (Figure 4B), activated in male oligodendrocytes, where the activated genes and pathways seem to indicate a process of motility and cell projection assembly, consistent with the processes observed in the previous section. Furthermore, another noteworthy point about this pathway is that its activation may originate from the VCAN gene secreted by neurons (yellow shadow), as can be seen in Figure 3H of the cell communication analysis. In the case of astrocytes (Figure 4C), the effectors differentially activated in males in the Axon guidance pathway lead mainly to axonal repulsion, such as the small GTPase RhoA, whose knockdown have been reported to promote true axon regeneration (PMCID: <u>PMC5436339</u>). Lastly, the AMPK signaling pathway in neurons (Figure 4D), which have been previously linked to neurodegenerative diseases (<u>PMID: 34355526</u>), is highly differentially activated in males, being EEF2 the principal deregulated effector, whose inactivation leads to protein synthesis inhibition and have been previously linked to the mechanism of action of LRRK2 in PD (<u>PMID: 32926469</u>). The rest of the pathways, together with the significant effector circuits by cell type, can be viewed in the associated web resource (https://bioinfo.cipf.es/cbl-atlas-pd/).

In the CT area, most significant subpathways appeared more activated in males, except for the microglia that showed more activated pathways in females (Figure S3A). Among these results, the numerous synapse pathways activated in male neurons or the presence of the apoptosis pathway in neurons are notable (Figure S3A). Only one apoptosis subpathway is significantly activated (TUBA1B effector), however when visualizing this pathway we see that many of the effectors show a tendency to be increased in females (Figure S3B).

### Comparison between substantia nigra and cortex meta-analyses

We next compared the meta-analyses results between brain regions in order to characterize sex differential expression profiles in brain regions involved in the disease. When we intersected the genetic signatures of each sex in each cell region for all cell types analyzed, neurons stood out as having a reversed pattern between sexes. That is, neurons in CT shared more genes with SN neurons of opposite sex than within the same sex (i.e., 26 common DEGs increased in SN male and in CT female, while no common DEGs between SN female and CT female) (Figure 5A). An exploratory analysis of protein-protein interaction in the genes of SN male and CT female intersection revealed a possible functional module mainly related to ribosomes, apoptosis, and the immune system. Notably this network obtained significantly more iterations than expected (PPI enrichment p-value: 2.49e-05). Due to their lower number, the DEGs common between SN female and CT male were characterised manually, with most of them related to synaptic processes, such as PHACTR3, involved in neurite outgrowth (PMID: <u>32423795</u>), or IQCJ-SCHIP1, implicated in action potential conduction and axon outgrowth (PMID: <u>25950943</u>).

**Figure 5.**
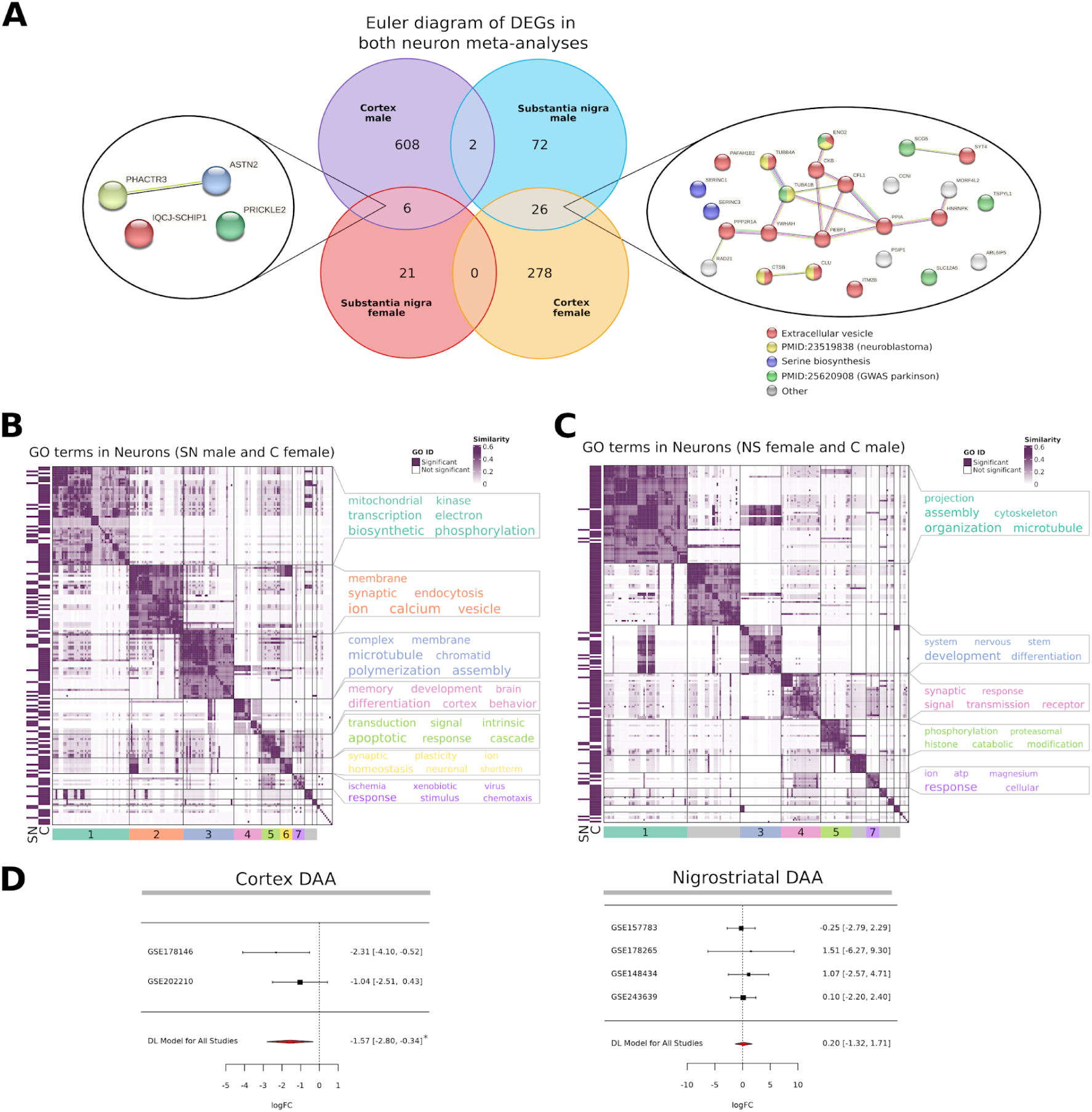
Comparison between the results obtained in the meta-analysis of the SN and CT circuits. A- Euler diagram with the neuron DEGs obtained in the meta-analysis of the nigrostriatal circuit and the cortex. Networks were obtained for intersections between opposite patterns, with an interaction confidence score threshold of 0.150. Through a literature search, we identify functions related to the genes and color them accordingly. B & C - Heatmap representation of the similarity matrix obtained from GO biological processes enriched in neurons in SN male and CT female meta-analyses (B) and SN female and CT male meta-analyses (C). Clusters are displayed at the bottom, and a wordcloud featuring the most significant terms within each cluster on the right side. The brain region to which each process belongs is illustrated on the left. D - Forest plot of differential abundance meta-analyses in CT and SN. The forest plots show the logFC and confidence intervals for each individual study and the DL model for all studies (lower red diamond). The asterisks indicate significant results (FDR < 0.05).

To further characterize this inverse pattern, we performed a semantic clustering of GO-BP functional signatures of the neurons from the two areas in opposite sexes. In the clustering between SN male and CT female (Figure 5B) all clusters identified presented a combination of the terms from both GO-BP sets, with no region-specific cluster obtained. The two largest clusters are related to the respiratory electron transport chain (cluster green) and to the calcium signaling (cluster orange), what can be related to a ER–mitochondria calcium crosstalk during the persistent stress that happens in PD, shifting the balance toward an apoptotic response (<u>Pizzo, P. et al. Cell Death Differ 2007</u>). This can be supported by the presence of functional clusters also related to apoptotic signaling pathway (cluster light green) and cellular response (cluster purple). On the other hand, in the clustering between SN female and CT male (Figure 5C) most common clusters are related with neuron system development (cluster blue) and synaptic signaling (cluster green and pink). However, unlike the previous one, this clustering showed some region-specific clusters (grey clusters), which indicates that not all biological processes identified are similar between these two regions.

To verify whether these biological processes were supported by differences in the number of neurons detected in each region, a meta-analysis of the DAAs done was also conducted (Figure 5D). The CT meta-analysis yielded a significant result indicating greater neurodegeneration in women with PD compared to their controls than in men, while in the SN meta-analysis, although significance was not reached, there was a slight opposite trend. However, these analyses may be biased by experimental and tissue variability across studies, which should be taken into account when interpreting these results.

## Discussion

With our meta-analyses of the SN and CT regions we evidence that there are differences between males and females at single cell transcriptome resolution level, with novel processes that deepen the understanding of sex differences in PD. Furthermore, our results are in accordance with the currently reported evidence about these differences (Cerri et al., 2019; DuMont et al., 2023; Nicoletti et al., 2023), including at the transcriptomic level (López-Cerdán et al., 2022). Our discussion will particularly emphasize the results obtained from the SN meta-analysis, since there are more studies from this region that support these results, and therefore, they are more robust than those obtained in the CT meta-analysis.

Neurons, the central cell type involved in PD, exhibit different phenotypes between sexes, with a phenotype indicative of heightened stress and predisposition to cell death in males, and a more connectivity preserved and neurotrophic supported one in females.

In male neurons, this phenotype is mainly characterized by proteostasis failure and metabolic dysregulation. This deficiency in proteostasis is notable in the dysregulation of genes related to alpha-synuclein processing and endosomal trafficking, highlighting the module comprising PRNP, PSAP, CLU, CTSB, and SORL1, most of them previously reported to be dysregulated in PD. The impairment of the former one is also potentially exacerbated by the interaction between the SORL1 receptor and APOE produced by oligodendrocytes. The upregulation of both SORL1 and APOE in myelinated neurons during early Parkinson’s disease (PD) pathogenesis has been previously reported (Wilhelmus et al., 2011), supporting the hypothesis that this receptor-ligand interaction may contribute to neurodegenerative mechanisms through disrupted intracellular trafficking and lipid metabolism. Furthermore, AMPK pathway and glycolytic impairment, although the literature is still ambiguous, can be an attempt of metabolic shifting amid chronic stress and mitochondrial dysfunction, with some of these glycolytic enzymes like ENO1/2 being previously described as markers of neuronal injury. These pathways have also been correlated with proteostasis failure (PMID: 34355526), which could directly lead to neuron apoptosis in male PD through ER stress, and necrotic features like axonal dystrophy (PMID: <u>31133790</u>).

On the counterpart, female neurons display a more homeostatic phenotype marked by enhanced neuronal differentiation, axonal maintenance, and dopaminergic resilience, showing better neurotrophic buffering and contributing to slower substantia nigra degeneration. This is further substantiated by the increase of the LMX1B gene, a critical transcription factor involved in the maintenance of dopaminergic neurons, whose inactivation has been shown to produce Parkinson’s-like cellular features (<u>PMC4968767</u>) and whose and in which SNPs positively associated with PD have been previously identified specifically in female subjects. (PMID: <u>19189040</u>). This resilience phenotype may respond to a potentiation of resilient neurons, probably due to estrogen effects (Y. H. Lee et al., 2019; Vegeto et al., 2019), although further studies are needed to test estrogen effects to harness these mechanisms.

This resilient neuronal phenotype is further bolstered by astrocyte supportive functions, particularly through enhanced astro-neuronal connectivity potentially mediated by GABBR1, a GABA-B receptor gene whose upregulation have been reported to enhance morphological complexity and connectivity with neurons (PMID: 39731735, 35373633, 28012274). This astrocytic phenotype can also be amplified by oligodendrocyte signaling via the SEMA5A-PLXNA2 axis, which would be expected to, by analogy to its role in neurons, regulate astrocyte morphology, process extension, and contact patterns through Rho-GTPase– dependent cytoskeletal changes (PMID: <u>34045951</u>). Complementing this, female oligodendrocytes exhibit a homeostatic, metabolically robust phenotype favoring myelin maintenance, with key genes like SPTLC2 (rate-limiting for sphingolipid/myelin lipid synthesis) and VPS4B (ATPase for endosomal membrane scission that influences cargo recycling and myelin membrane turnover). Since myelination and lipid dysregulation is implicated in PD and white-matter pathology (PMCID: <u>PMC9303520</u>), this myelinating phenotype can be a female-specific counteract myelin decline to preserve neuronal function during the disease.

The neuroinflammatory response is highly relevant in the development of PD, contributing directly and indirectly to oxidative stress and neuronal degeneration (Boyd et al., 2022; Mukhara et al., 2020). Together with astrocytes, microglia are the main cells involved in this process, and can adopt both detrimental and protective phenotypes in response to different stimuli (Terrin et al., 2023). In our results, male microglia displayed a predominantly pro-inflammatory phenotype, characterized by genes such as FCGR2A, an Fcγ receptor that drives phagocytosis and amplifies immune signaling, and LAPTM5, a lysosomal adaptor linking endocytosis to cytokine production. These pathways suggest a strong activation of antigen presentation and inflammatory cascades, which may exacerbate neuronal stress and accelerate neurodegeneration in PD (Bartels et al., 2020). In contrast, female microglia showed a profile more oriented toward clearance and homeostatic regulation, with enrichment for genes involved in phagosome maturation and autophagy, including PICALM and SORL1, as well as regulatory elements such as TGFBR2 and RGS10, both previously associated with protective microglial states. This functional bias toward aggregate clearance and controlled immune activation could mitigate proteostatic burden and limit chronic inflammation, potentially contributing to sex-related differences in PD progression. Notably, prior literature postulates the effect of sex hormones as a possible etiology of the lower neuroinflammation in females (Cerri et al., 2019; Terrin et al., 2023; Yanguas-Casás, 2020). These hormonal influences may underlie the observed tendency of female microglia to maintain homeostasis and efficient clearance mechanisms, highlighting a possible link between endocrine factors and resilience to neurodegenerative stress in PD.

The role of protein aggregate clearance in PD appears to be strongly influenced by sex-specific microglial function. Processes related to amyloid-beta metabolism and clearance are enriched across multiple cell types in males—neurons, astrocytes, and oligodendrocytes— whereas in females these processes are largely restricted to microglia (Figure 3G). This pattern suggests that male microglia may exhibit impaired capacity for aggregate processing, forcing other cell types to activate protein aggregates reaction pathways. While these responses aim to mitigate proteotoxic stress, they likely impose a significant metabolic burden, promoting cellular stress and inflammatory signaling in neurons and glia. Such stress responses can, in turn, amplify crosstalk with microglia, reinforcing pro-inflammatory cascades and creating a self-perpetuating loop of cellular stress and immune activation that may accelerate neurodegeneration in males.

Previous studies with transcriptomic-based approaches and sex perspective have also related males with a phenotype more related to the pathological processes associated with PD (López-Cerdán et al., 2022; Simunovic et al., 2010). However, these sex-associated patterns of neurodegeneration and synapse potentiation are inverted in the cortical region results. In males, our results suggest a higher neuron connectivity in the cortical area, which have been described especially in patients with rigidity-dominant PD, also more common in males (Hou et al., 2017, 2018; Wichmann, 2019). We also observed in males processes related with a greater loss of neurons in the nigrostriatal system, existing bibliography that relate this to freezing of gait symptom, which is also more common in males (Bohnen et al., 2019; Kim et al., 2018). In contrast, our results suggest more neurodegeneration in cortical areas and a milder nigrostriatal impairment in female patients, being the former related to the tremor-dominant PD, more common among females (Haaxma et al., 2007). Furthermore, some studies have related dysfunction in some areas of the frontal cortex and a reduced anterior cingulate cortex with anxiety and depression states, also predominant symptoms in female PD patients (Carey et al., 2021; Laux, 2022). All this may indicate sex differences not only in the molecular mechanisms involved in disease progression but also in brain regions differentially affected by PD, with characteristic clinical implications.

In conclusion, we have identified different biological processes that outline a heightened immune response in males across both C and NS regions, and a greater neurodegenerative process in the NS in males compared to females, which seems to be inverted in the cortical region. Given the various approaches used, we identified signaling pathways, genes and cellular interactions involved in these differences, which may provide new insights to understand the molecular mechanisms of PD that can help to improve its diagnosis and personalized treatment.

We obtained these results through an *in silico* analysis of data available in public repositories, which demonstrates the power of using this type of approach to obtain more insight into the etiopathogenesis of different diseases from previously generated data. To promote these approaches, it is essential to conduct open research and make data available in public repositories according to FAIR principles (Findable, Accessible, Interoperable, Reusable), with full information on covariates of interest, such as sex. In this regard, a major pitfall in this meta-analysis was the inclusion of eligible studies with equal or similar representation of both sexes in each experimental condition. The most common scenario is the underrepresentation of females in this type of studies, which has been widely reported (Vaidya et al., 2021; Vitale et al., 2017; Weisman & Cassard, 1999). In our case, a representative example is the GSE157783 study, with only 1 female sample in each condition. This underrepresentation not only hinders *in silico* approaches as in our work, but can also introduce biases in the original studies, making it difficult to extrapolate the results to general society. We consider it necessary to ensure the inclusion of sex information in the studies performed as well as the inclusion of an appropriate number of each of the sexes in the conditions analyzed.

We also acknowledge that C meta-analysis may present less robustness than NS meta-analysis, since it is composed of only 2 studies. Moreover, each study comes from a different cortical region (anterior cingulate cortex and prefrontal cortex), nevertheless, these regions are physically very close and closely related in the progression of the disease. (Braak et al., 2003; Del Tredici & Braak, 2016).

We tried to avoid these limitations by using a meta-analysis approach, an integration strategy that has never been used before in snRNA-seq datasets. This approach allows us to avoid biases arising from each individual study, and to work with relatively smaller datasets than if we tried to integrate all the data in one large dataset, increasing the reliability of the results and reducing the time and computational cost of the analyses. In addition, selecting the major cell types to perform the analyses allows us to work with a large number of cells from all the patients, which also increases the robustness of the results obtained. Finally, although sex-based meta-analyses of transcriptomic studies in this disease have been performed previously (López-Cerdán et al., 2022; Mariani et al., 2018), performing them on single cell technologies provides greater specificity and clarity of the results obtained. This could help to better interpret possible sex differences in PD by exploring the processes involved in each cell type, as well as the possible interactions between them.

In conclusion, this work has identified sex differences in potential genes, biological processes and pathways relevant to the pathogenesis of PD at cell type resolution. These results may help to better understand the clinical differences found between females and males and potentiate sex-based targeted therapy or drug repositioning research. As a final statement, we emphasize the importance of sharing data on open access platforms to facilitate scientific advancement.

## Bibliography

Alexa, A., & Rahnenfuhrer, J. (2023). topGO: Enrichment Analysis for Gene Ontology (Version R package version 2.54.0) [Computer software]. https://bioconductor.org/packages/topGO

Alexa, A., Rahnenführer, J., & Lengauer, T. (2006). Improved scoring of functional groups from gene expression data by decorrelating GO graph structure. Bioinformatics, 22(13), 1600–1607. 10.1093/bioinformatics/btl140

Amezquita, R. A., Lun, A. T. L., Becht, E., Carey, V. J., Carpp, L. N., Geistlinger, L., Marini, F., Rue-Albrecht, K., Risso, D., Soneson, C., Waldron, L., Pagès, H., Smith, M. L., Huber, W., Morgan, M., Gottardo, R., & Hicks, S. C. (2020). Orchestrating Single-Cell Analysis with Bioconductor. Nature Methods, 17(2), 137–145. 10.1038/s41592-019-0654-x

Aran, D., Looney, A. P., Liu, L., Wu, E., Fong, V., Hsu, A., Chak, S., Naikawadi, R. P., Wolters, P. J., Abate, A. R., Butte, A. J., & Bhattacharya, M. (2019). Reference-based analysis of lung single-cell sequencing reveals a transitional profibrotic macrophage. Nature Immunology, 20(2), Article 2. 10.1038/s41590-018-0276-y

Ascherio, A., LeWitt, P. A., Xu, K., Eberly, S., Watts, A., Matson, W. R., Marras, C., Kieburtz, K., Rudolph, A., Bogdanov, M. B., Schwid, S. R., Tennis, M., Tanner, C. M., Beal, M. F., Lang, A. E., Oakes, D., Fahn, S., Shoulson, I., Schwarzschild, M. A., & Parkinson Study Group DATATOP Investigators. (2009). Urate as a Predictor of the Rate of Clinical Decline in Parkinson Disease. Archives of Neurology, 66(12), 1460–1468. 10.1001/archneurol.2009.247

Bajo-Grañeras, R., Ganfornina, M. D., Martín-Tejedor, E., & Sanchez, D. (2011). Apolipoprotein D mediates autocrine protection of astrocytes and controls their reactivity level, contributing to the functional maintenance of paraquat-challenged dopaminergic systems. Glia, 59(10), 1551–1566. 10.1002/glia.21200

Bakken, T. E., Jorstad, N. L., Hu, Q., Lake, B. B., Tian, W., Kalmbach, B. E., Crow, M., Hodge, R. D., Krienen, F. M., Sorensen, S. A., Eggermont, J., Yao, Z., Aevermann, B. D., Aldridge, A. I., Bartlett, A., Bertagnolli, D., Casper, T., Castanon, R. G., Crichton, K., … Lein, E. S. (2021). Allen Institute for Brain Science (2020). Allen Cell Types Database—Human M1—10x Genomics [dataset]. Available from celltypes.brain-map.org/rnaseq. Primary publication: Comparative cellular analysis of motor cortex in human, marmoset and mouse. Nature, 598(7879), Article 7879. 10.1038/s41586-021-03465-8

Barko, K., Shelton, M., Xue, X., Afriyie-Agyemang, Y., Puig, S., Freyberg, Z., Tseng, G. C., Logan, R. W., & Seney, M. L. (2022). Brain region- and sex-specific transcriptional profiles of microglia. Frontiers in Psychiatry, 13. https://www.frontiersin.org/journals/psychiatry/articles/10.3389/fpsyt.2022.945548

Bartels, T., De Schepper, S., & Hong, S. (2020). Microglia modulate neurodegeneration in Alzheimer’s and Parkinson’s diseases. Science, 370(6512), 66–69. 10.1126/science.abb8587

Belvisi, D., Pellicciari, R., Fabbrini, G., Tinazzi, M., Berardelli, A., & Defazio, G. (2020). Modifiable risk and protective factors in disease development, progression and clinical subtypes of Parkinson’s disease: What do prospective studies suggest? Neurobiology of Disease, 134, 104671. 10.1016/j.nbd.2019.104671

Blighe, K., & Lun, A. (2021). PCAtools: Everything Principal Components Analysis. (Version 2.6.0.) [Computer software]. https://github.com/kevinblighe/PCAtools

Bloem, B. R., Okun, M. S., & Klein, C. (2021). Parkinson’s disease. Lancet (London, England), 397(10291), 2284–2303. 10.1016/S0140-6736(21)00218-X

Bohnen, N. I., Kanel, P., Zhou, Z., Koeppe, R. A., Frey, K. A., Dauer, W. T., Albin, R. L., & Müller, M. L. T. M. (2019). Cholinergic system changes of falls and freezing of gait in Parkinson’s disease. Annals of Neurology, 85(4), 538–549. 10.1002/ana.25430

Bourque, M., Morissette, M., Soulet, D., & Di Paolo, T. (2023). Impact of sex on neuroimmune contributions to Parkinson’s disease. Brain Research Bulletin, 199, 110668. 10.1016/j.brainresbull.2023.110668

Boyd, R. J., Avramopoulos, D., Jantzie, L. L., & McCallion, A. S. (2022). Neuroinflammation represents a common theme amongst genetic and environmental risk factors for Alzheimer and Parkinson diseases. Journal of Neuroinflammation, 19(1), 223. 10.1186/s12974-022-02584-x

Braak, H., Tredici, K. D., Rüb, U., de Vos, R. A. I., Jansen Steur, E. N. H., & Braak, E. (2003). Staging of brain pathology related to sporadic Parkinson’s disease. Neurobiology of Aging, 24(2), 197–211. 10.1016/S0197-4580(02)00065-9

Cabrera, J. R., Bouzas-Rodriguez, J., Tauszig-Delamasure, S., & Mehlen, P. (2011). RET Modulates Cell Adhesion via Its Cleavage by Caspase in Sympathetic Neurons *. Journal of Biological Chemistry, 286(16), 14628–14638. 10.1074/jbc.M110.195461

Carey, G., Görmezoğlu, M., de Jong, J. J. A., Hofman, P. A. M., Backes, W. H., Dujardin, K., & Leentjens, A. F. G. (2021). Neuroimaging of Anxiety in Parkinson’s Disease: A Systematic Review. Movement Disorders, 36(2), 327–339. 10.1002/mds.28404

Cerri, S., Mus, L., & Blandini, F. (2019). Parkinson’s Disease in Women and Men: What’s the Difference? Journal of Parkinson’s Disease, 9(3), 501–515. 10.3233/JPD-191683

Chaudry, O., Ndukwe, K., Xie, L., Figueiredo-Pereira, M., Serrano, P., & Rockwell, P. (2022). Females exhibit higher GluA2 levels and outperform males in active place avoidance despite increased amyloid plaques in TgF344-Alzheimer’s rats. Scientific Reports, 12. 10.1038/s41598-022-23801-w

de Kanter, J. K., Lijnzaad, P., Candelli, T., Margaritis, T., & Holstege, F. C. P. (2019). CHETAH: A selective, hierarchical cell type identification method for single-cell RNA sequencing. Nucleic Acids Research, 47(16), e95. 10.1093/nar/gkz543

Del Tredici, K., & Braak, H. (2016). Review: Sporadic Parkinson’s disease: development and distribution of α-synuclein pathology. Neuropathology and Applied Neurobiology, 42(1), 33–50. 10.1111/nan.12298

Dong-Chen, X., Yong, C., Yang, X., Chen-Yu, S., & Li-Hua, P. (2023). Signaling pathways in Parkinson’s disease: Molecular mechanisms and therapeutic interventions. Signal Transduction and Targeted Therapy, 8, 73. 10.1038/s41392-023-01353-3

Drinkut, A., Tillack, K., Meka, D. P., Schulz, J. B., Kügler, S., & Kramer, E. R. (2016). Ret is essential to mediate GDNF’s neuroprotective and neuroregenerative effect in a Parkinson disease mouse model. Cell Death & Disease, 7(9), e2359–e2359. 10.1038/cddis.2016.263

DuMont, M., Agostinis, A., Singh, K., Swan, E., Buttle, Y., & Tropea, D. (2023). Sex representation in neurodegenerative and psychiatric disorders’ preclinical and clinical studies. Neurobiology of Disease, 184, 106214. 10.1016/j.nbd.2023.106214

Feleke, R., Reynolds, R. H., Smith, A. M., Tilley, B., Taliun, S. A. G., Hardy, J., Matthews, P. M., Gentleman, S., Owen, D. R., Johnson, M. R., Srivastava, P. K., & Ryten, M. (2021). Cross-platform transcriptional profiling identifies common and distinct molecular pathologies in Lewy body diseases. Acta Neuropathologica, 142(3), 449–474. 10.1007/s00401-021-02343-x

Flores-Cuadrado, A., Saiz-Sanchez, D., Mohedano-Moriano, A., Lamas-Cenjor, E., Leon-Olmo, V., Martinez-Marcos, A., & Ubeda-Bañon, I. (2021). Astrogliosis and sexually dimorphic neurodegeneration and microgliosis in the olfactory bulb in Parkinson’s disease. NPJ Parkinson’s Disease, 7, 11. 10.1038/s41531-020-00154-7

Gray, A. L., Hyde, T. M., Deep-Soboslay, A., Kleinman, J. E., & Sodhi, M. S. (2015). Sex differences in glutamate receptor gene expression in major depression and suicide. Molecular Psychiatry, 20(9), Article 9. 10.1038/mp.2015.91

Haaxma, C. A., Bloem, B. R., Borm, G. F., Oyen, W. J. G., Leenders, K. L., Eshuis, S., Booij, J., Dluzen, D. E., & Horstink, M. W. I. M. (2007). Gender differences in Parkinson’s disease. *Journal of Neurology*, Neurosurgery & Psychiatry, 78(8), 819–824. 10.1136/jnnp.2006.103788

Hidalgo, M. R., Cubuk, C., Amadoz, A., Salavert, F., Carbonell-Caballero, J., & Dopazo, J. (2016). High throughput estimation of functional cell activities reveals disease mechanisms and predicts relevant clinical outcomes. Oncotarget, 8(3), 5160–5178. 10.18632/oncotarget.14107

Hinarejos, I., Machuca, C., Sancho, P., & Espinós, C. (2020). Mitochondrial Dysfunction, Oxidative Stress and Neuroinflammation in Neurodegeneration with Brain Iron Accumulation (NBIA). Antioxidants, 9(10). 10.3390/antiox9101020

Hong, G. P., Kim, M.-H., & Kim, H. J. (2021). Sex-related Differences in Glial Fibrillary Acidic Protein-positive GABA Regulate Neuropathology Following Pilocarpine-induced Status Epilepticus. Neuroscience, 472, 157–166. 10.1016/j.neuroscience.2021.08.002

Hou, Y., Luo, C., Yang, J., Ou, R., Liu, W., Song, W., Gong, Q., & Shang, H. (2017). Default-mode network connectivity in cognitively unimpaired drug-naïve patients with rigidity-dominant Parkinson’s disease. Journal of Neurology, 264(1), 152–160. 10.1007/s00415-016-8331-9

Hou, Y., Wei, Q., Ou, R., Yang, J., Song, W., Gong, Q., & Shang, H. (2018). Impaired topographic organization in cognitively unimpaired drug-naïve patients with rigidity-dominant Parkinson’s disease. Parkinsonism & Related Disorders, 56, 52–57. 10.1016/j.parkreldis.2018.06.021

Iovino, L., Tremblay, M. E., & Civiero, L. (2020). Glutamate-induced excitotoxicity in Parkinson’s disease: The role of glial cells. Journal of Pharmacological Sciences, 144(3), 151–164. 10.1016/j.jphs.2020.07.011

Junqueira, S. C., Centeno, E. G. Z., Wilkinson, K. A., & Cimarosti, H. (2019). Post-translational modifications of Parkinson’s disease-related proteins: Phosphorylation, SUMOylation and Ubiquitination. Biochimica et Biophysica Acta (BBA) - Molecular Basis of Disease, 1865(8), 2001–2007. 10.1016/j.bbadis.2018.10.025

Kamath, T., Abdulraouf, A., Burris, S. J., Gazestani, V., Nadaf, N., Vanderburg, C., & Macosko, E. Z. (2021). A molecular census of midbrain dopaminergic neurons in Parkinson’s disease (p. 2021.06.16.448661). bioRxiv. 10.1101/2021.06.16.448661

Kim, R., Lee, J., Kim, Y., Kim, A., Jang, M., Kim, H.-J., Jeon, B., Kang, U. J., & Fahn, S. (2018). Presynaptic striatal dopaminergic depletion predicts the later development of freezing of gait in de novo Parkinson’s disease: An analysis of the PPMI cohort. Parkinsonism & Related Disorders, 51, 49–54. 10.1016/j.parkreldis.2018.02.047

Lane, R. F., Raines, S. M., Steele, J. W., Ehrlich, M. E., Lah, J. A., Small, S. A., Tanzi, R. E., Attie, A. D., & Gandy, S. (2010). Diabetes-Associated SorCS1 Regulates Alzheimer’s Amyloid-β Metabolism: Evidence for Involvement of SorL1 and the Retromer Complex. Journal of Neuroscience, 30(39), 13110–13115. 10.1523/JNEUROSCI.3872-10.2010

Laux, G. (2022). Parkinson and depression: Review and outlook. Journal of Neural Transmission, 129(5), 601–608. 10.1007/s00702-021-02456-3

Lee, A. J., Kim, C., Park, S., Jun, K., Eom, J., Lee, S.-J., Chung, S. J., Rissman, R. A., Chung, J., Masliah, E., & Jung, I. (2022). *Characterization of altered molecular mechanisms in Parkinson’s disease* through *cell type-resolved multi-omics analyses* (p. 2022.02.13.479386). bioRxiv. 10.1101/2022.02.13.479386

Lee, Y. H., Cha, J., Chung, S. J., Yoo, H. S., Sohn, Y. H., Ye, B. S., & Lee, P. H. (2019). Beneficial effect of estrogen on nigrostriatal dopaminergic neurons in drug-naïve postmenopausal Parkinson’s disease. Scientific Reports, 9(1), Article 1. 10.1038/s41598-019-47026-6

Liberati, A., Altman, D. G., Tetzlaff, J., Mulrow, C., Gøtzsche, P. C., Ioannidis, J. P. A., Clarke, M., Devereaux, P. J., Kleijnen, J., & Moher, D. (2009). The PRISMA statement for reporting systematic reviews and meta-analyses of studies that evaluate healthcare interventions: Explanation and elaboration. The BMJ, 339, b2700. 10.1136/bmj.b2700

Liu, Y., Shen, X., Zhang, Y., Zheng, X., Cepeda, C., Wang, Y., Duan, S., & Tong, X. (2023). Interactions of glial cells with neuronal synapses, from astrocytes to microglia and oligodendrocyte lineage cells. Glia, 71(6), 1383–1401. 10.1002/glia.24343

López-Cerdán, A., Andreu, Z., Hidalgo, M. R., Grillo-Risco, R., Català-Senent, J. F., Soler-Sáez, I., Neva-Alejo, A., Gordillo, F., de la Iglesia-Vayá, M., & García-García, F. (2022). Unveiling sex-based differences in Parkinson’s disease: A comprehensive meta-analysis of transcriptomic studies. Biology of Sex Differences, 13, 68. 10.1186/s13293-022-00477-5

Lotharius, J., & Brundin, P. (2002). Pathogenesis of parkinson’s disease: Dopamine, vesicles and α-synuclein. Nature Reviews Neuroscience, 3(12), Article 12. 10.1038/nrn983

Luca, A., Monastero, R., Cicero, C. E., Baschi, R., Donzuso, G., Mostile, G., Restivo, V., Di Giorgi, L., Caccamo, M., Zappia, M., & Nicoletti, A. (2022). Executive functioning and serum lipid fractions in Parkinson’s disease—a possible sex-effect: The PACOS study. Journal of Neural Transmission, 129(3), 287–293. 10.1007/s00702-022-02460-1

Lun, A. T. L., McCarthy, D. J., & Marioni, J. C. (2016). *A step-by-step workflow for low-level analysis of single-cell RNA-seq data with Bioconductor* (5:2122). F1000Research. 10.12688/f1000research.9501.2

Lun, A. T. L., Riesenfeld, S., Andrews, T., Dao, T. P., Gomes, T., Marioni, J. C., & participants in the 1st Human Cell Atlas Jamboree. (2019). EmptyDrops: Distinguishing cells from empty droplets in droplet-based single-cell RNA sequencing data. Genome Biology, 20(1), 63. 10.1186/s13059-019-1662-y

Maiti, P., Manna, J., & Dunbar, G. L. (2017). Current understanding of the molecular mechanisms in Parkinson’s disease: Targets for potential treatments. Translational Neurodegeneration, 6(1), 28. 10.1186/s40035-017-0099-z

Mariani, E., Lombardini, L., Facchin, F., Pizzetti, F., Frabetti, F., Tarozzi, A., & Casadei, R. (2018). Sex-Specific Transcriptome Differences in Substantia Nigra Tissue: A Meta-Analysis of Parkinson’s Disease Data. Genes, 9(6), 275. 10.3390/genes9060275

McDavid, A., Finak, G., & Yajima, M. (2021). MAST: Model-based Analysis of Single Cell Transcriptomics. (Version 1.20.0.) [Computer software]. https://github.com/RGLab/MAST/

McKenzie, A., Wang, M., & Zhang, B. (2021). BRETIGEA: Brain Cell Type Specific Gene Expression Analysis. (Version 1.0.3.) [Computer software]. https://CRAN.R-project.org/package=BRETIGEA

Meoni, S., Macerollo, A., & Moro, E. (2020). Sex differences in movement disorders. Nature Reviews Neurology, 16(2), Article 2. 10.1038/s41582-019-0294-x

Miyazaki, I., Asanuma, M., Kikkawa, Y., Takeshima, M., Murakami, S., Miyoshi, K., Sogawa, N., & Kita, T. (2011). Astrocyte-derived metallothionein protects dopaminergic neurons from dopamine quinone toxicity. Glia, 59(3), 435–451. 10.1002/glia.21112

Mukhara, D., Oh, U., & Neigh, G. N. (2020). Neuroinflammation. Handbook of Clinical Neurology, 175, 235–259. 10.1016/B978-0-444-64123-6.00017-5

Najyb, O., Do Carmo, S., Alikashani, A., & Rassart, E. (2017). Apolipoprotein D Overexpression Protects Against Kainate-Induced Neurotoxicity in Mice. Molecular Neurobiology, 54(6), 3948–3963. 10.1007/s12035-016-9920-4

Nicoletti, A., Baschi, R., Cicero, C. E., Iacono, S., Re, V. L., Luca, A., Schirò, G., Monastero, R., & Gender Neurology Study Group of the Italian Society of Neurology. (2023). Sex and gender differences in Alzheimer’s disease, Parkinson’s disease, and Amyotrophic Lateral Sclerosis: A narrative review. Mechanisms of Ageing and Development, 212, 111821. 10.1016/j.mad.2023.111821

Page, M. J., McKenzie, J. E., Bossuyt, P. M., Boutron, I., Hoffmann, T. C., Mulrow, C. D., Shamseer, L., Tetzlaff, J. M., Akl, E. A., Brennan, S. E., Chou, R., Glanville, J., Grimshaw, J. M., Hróbjartsson, A., Lalu, M. M., Li, T., Loder, E. W., Mayo-Wilson, E., McDonald, S., … Moher, D. (2021). The PRISMA 2020 statement: An updated guideline for reporting systematic reviews. Journal of Clinical Epidemiology, 134, 178–189. 10.1016/j.jclinepi.2021.03.001

Panicker, N., Ge, P., Dawson, V. L., & Dawson, T. M. (2021). The cell biology of Parkinson’s disease. The Journal of Cell Biology, 220(4), e202012095. 10.1083/jcb.202012095

Patani, R., Hardingham, G. E., & Liddelow, S. A. (2023). Functional roles of reactive astrocytes in neuroinflammation and neurodegeneration. Nature Reviews Neurology, 19(7), Article 7. 10.1038/s41582-023-00822-1

Picillo, M., Nicoletti, A., Fetoni, V., Garavaglia, B., Barone, P., & Pellecchia, M. T. (2017). The relevance of gender in Parkinson’s disease: A review. Journal of Neurology, 264(8), 1583–1607. 10.1007/s00415-016-8384-9

R Core Team. (2021). *R: A language and environment for statistical computing*. (Version 4.1.2) [Computer software]. R Foundation for Statistical Computing. https://www.R-project.org/

Raghupathy, R., Al-Mutawa, E., Al-Azemi, M., Makhseed, M., Azizieh, F., & Szekeres-Bartho, J. (2009). Progesterone-induced blocking factor (PIBF) modulates cytokine production by lymphocytes from women with recurrent miscarriage or preterm delivery. Journal of Reproductive Immunology, 80(1), 91–99. 10.1016/j.jri.2009.01.004

Simon, D. K., Tanner, C. M., & Brundin, P. (2020). Parkinson Disease Epidemiology, Pathology, Genetics and Pathophysiology. Clinics in Geriatric Medicine, 36(1), 1–12. 10.1016/j.cger.2019.08.002

Simunovic, F., Yi, M., Wang, Y., Stephens, R., & Sonntag, K. C. (2010). Evidence for Gender-Specific Transcriptional Profiles of Nigral Dopamine Neurons in Parkinson Disease. PLoS ONE, 5(1), e8856. 10.1371/journal.pone.0008856

Smajić, S., Prada-Medina, C. A., Landoulsi, Z., Ghelfi, J., Delcambre, S., Dietrich, C., Jarazo, J., Henck, J., Balachandran, S., Pachchek, S., Morris, C. M., Antony, P., Timmermann, B., Sauer, S., Pereira, S. L., Schwamborn, J. C., May, P., Grünewald, A., & Spielmann, M. (2021). Single-cell sequencing of human midbrain reveals glial activation and a Parkinson-specific neuronal state. Brain, 145(3), 964–978. 10.1093/brain/awab446

Snel, B., Lehmann, G., Bork, P., & Huynen, M. A. (2000). STRING: A web-server to retrieve and display the repeatedly occurring neighbourhood of a gene. Nucleic Acids Research, 28(18), 3442–3444.

Szekeres-Bartho, J., Barakonyi, A., Par, G., Polgar, B., Palkovics, T., & Szereday, L. (2001). Progesterone as an immunomodulatory molecule. International Immunopharmacology, 1(6), 1037– 1048. 10.1016/S1567-5769(01)00035-2

Szklarczyk, D., Gable, A. L., Nastou, K. C., Lyon, D., Kirsch, R., Pyysalo, S., Doncheva, N. T., Legeay, M., Fang, T., Bork, P., Jensen, L. J., & von Mering, C. (2020). The STRING database in 2021: Customizable protein–protein networks, and functional characterization of user-uploaded gene/measurement sets. Nucleic Acids Research, 49(D1), D605–D612. 10.1093/nar/gkaa1074

Terrin, F., Tesoriere, A., Plotegher, N., & Dalla Valle, L. (2023). Sex and Brain: The Role of Sex Chromosomes and Hormones in Brain Development and Parkinson’s Disease. Cells, 12(11), 1486. 10.3390/cells12111486

Tolosa, E., Garrido, A., Scholz, S. W., & Poewe, W. (2021). Challenges in the diagnosis of Parkinson’s disease. The Lancet. Neurology, 20(5), 385–397. 10.1016/S1474-4422(21)00030-2

Tsika, E., Glauser, L., Moser, R., Fiser, A., Daniel, G., Sheerin, U.-M., Lees, A., Troncoso, J. C., Lewis, P. A., Bandopadhyay, R., Schneider, B. L., & Moore, D. J. (2014). Parkinson’s disease-linked mutations in VPS35 induce dopaminergic neurodegeneration. Human Molecular Genetics, 23(17), 4621–4638. 10.1093/hmg/ddu178

Türei, D., Valdeolivas, A., Gul, L., Palacio-Escat, N., Klein, M., Ivanova, O., Ölbei, M., Gábor, A., Theis, F., Módos, D., Korcsmáros, T., & Saez-Rodriguez, J. (2021). Integrated intra- and intercellular signaling knowledge for multicellular omics analysis. Molecular Systems Biology, 17(3), e9923. 10.15252/msb.20209923

Vaidya, B., Dhamija, K., Guru, P., & Sharma, S. S. (2021). Parkinson’s disease in women: Mechanisms underlying sex differences. European Journal of Pharmacology, 895, 173862. 10.1016/j.ejphar.2021.173862

Vázquez-Vélez, G. E., & Zoghbi, H. Y. (2021). Parkinson’s Disease Genetics and Pathophysiology. Annual Review of Neuroscience, 44, 87–108. 10.1146/annurev-neuro-100720-034518

Vegeto, E., Villa, A., Della Torre, S., Crippa, V., Rusmini, P., Cristofani, R., Galbiati, M., Maggi, A., & Poletti, A. (2019). The Role of Sex and Sex Hormones in Neurodegenerative Diseases. Endocrine Reviews, 41(2), 273–319. 10.1210/endrev/bnz005

Verma, M., Lizama, B. N., & Chu, C. T. (2022). Excitotoxicity, calcium and mitochondria: A triad in synaptic neurodegeneration. Translational Neurodegeneration, 11(1), 3. 10.1186/s40035-021-00278-7

Viechtbauer, W. (2010). Conducting Meta-Analyses in R with the metafor Package. Journal of Statistical Software, 36, 1–48. 10.18637/jss.v036.i03

Villa, A., Gelosa, P., Castiglioni, L., Cimino, M., Rizzi, N., Pepe, G., Lolli, F., Marcello, E., Sironi, L., Vegeto, E., & Maggi, A. (2018). Sex-Specific Features of Microglia from Adult Mice. Cell Reports, 23(12), 3501–3511. 10.1016/j.celrep.2018.05.048

Vitale, C., Fini, M., Spoletini, I., Lainscak, M., Seferovic, P., & Rosano, G. M. (2017). Under-representation of elderly and women in clinical trials. International Journal of Cardiology, 232, 216–221. 10.1016/j.ijcard.2017.01.018

Walden, H., & Muqit, M. M. K. (2017). Ubiquitin and Parkinson’s disease through the looking glass of genetics. Biochemical Journal, 474(9), 1439–1451. 10.1042/BCJ20160498

Wang, Q., Wang, M., Choi, I., Sarrafha, L., Liang, M., Ho, L., Farrell, K., Beaumont, K. G., Sebra, R., De Sanctis, C., Crary, J. F., Ahfeldt, T., Blanchard, J., Neavin, D., Powell, J., Davis, D. A., Sun, X., Zhang, B., & Yue, Z. (2024). Molecular profiling of human substantia nigra identifies diverse neuron types associated with vulnerability in Parkinson’s disease. Science Advances, 10(2), eadi8287. 10.1126/sciadv.adi8287

Wang, Y., Wang, R., Zhang, S., Song, S., Jiang, C., Han, G., Wang, M., Ajani, J., Futreal, A., & Wang, L. (2019). iTALK: An R Package to Characterize and Illustrate Intercellular Communication (p. 507871). bioRxiv. 10.1101/507871

Wang, Z.-L., Yuan, L., Li, W., & Li, J.-Y. (2022). Ferroptosis in Parkinson’s disease: Glia–neuron crosstalk. Trends in Molecular Medicine, 28(4), 258–269. 10.1016/j.molmed.2022.02.003

Weisman, C. S., & Cassard, S. D. (1999). Health Consequences of Exclusion or Underrepresentation of Women in Clinical Studies (I). In Women and Health Research: Ethical and Legal Issues of Including Women in Clinical Studies: Volume 2: Workshop and Commissioned Papers. National Academies Press (US). https://www.ncbi.nlm.nih.gov/books/NBK236583/

Wichmann, T. (2019). Changing views of the pathophysiology of Parkinsonism. Movement Disorders, 34(8), 1130–1143. 10.1002/mds.27741

Wilhelmus, M. M. M., Bol, J. G. J. M., Haastert, E. S. V., Rozemuller, A. J. M., Bu, G., Drukarch, B., & Hoozemans, J. J. M. (2011). Apolipoprotein E and LRP1 Increase Early in Parkinson’s Disease Pathogenesis. The American Journal of Pathology, 179(5), 2152–2156. 10.1016/j.ajpath.2011.07.021

Xu, J., Farsad, H. L., Hou, Y., Barclay, K., Lopez, B. A., Yamada, S., Saliu, I. O., Shi, Y., Knight, W. C., Bateman, R. J., Benzinger, T. L. S., Yi, J. J., Li, Q., Wang, T., Perlmutter, J. S., Morris, J. C., & Zhao, G. (2023). Human striatal glia differentially contribute to AD- and PD-specific neurodegeneration. Nature Aging, 3(3), Article 3. 10.1038/s43587-023-00363-8

Yin, J., Spillman, E., Cheng, E. S., Short, J., Chen, Y., Lei, J., Gibbs, M., Rosenthal, J. S., Sheng, C., Chen, Y. X., Veerasammy, K., Choetso, T., Abzalimov, R., Wang, B., Han, C., He, Y., & Yuan, Q. (2021). Brain-specific lipoprotein receptors interact with astrocyte derived apolipoprotein and mediate neuron-glia lipid shuttling. Nature Communications, 12, 2408. 10.1038/s41467-021-22751-7

Zecca, L., Youdim, M. B. H., Riederer, P., Connor, J. R., & Crichton, R. R. (2004). Iron, brain ageing and neurodegenerative disorders. Nature Reviews Neuroscience, 5(11), Article 11. 10.1038/nrn1537

Zhang, N., Yu, X., Song, L., Xiao, Z., Xie, J., & Xu, H. (2022). Ferritin confers protection against iron-mediated neurotoxicity and ferroptosis through iron chelating mechanisms in MPP+-induced MES23.5 dopaminergic cells. Free Radical Biology and Medicine, 193, 751–763. 10.1016/j.freeradbiomed.2022.11.018

Zhang, X., Lan, Y., Xu, J., Quan, F., Zhao, E., Deng, C., Luo, T., Xu, L., Liao, G., Yan, M., Ping, Y., Li, F., Shi, A., Bai, J., Zhao, T., Li, X., & Xiao, Y. (2019). CellMarker: A manually curated resource of cell markers in human and mouse. Nucleic Acids Research, 47(Database issue), D721–D728. 10.1093/nar/gky900

Zhang, Z. (2019). SCINA: A Semi-Supervised Category Identification and Assignment Tool. (Version 1.2.0.) [Computer software]. https://CRAN.R-project.org/package=SCINA

Zhu, B., Park, J.-M., Coffey, S., Hsu, I.-U., Lam, T. T., Gopal, P. P., Ginsberg, S. D., Wang, J., Su, C., Zhao, H., Hafler, D. A., Chandra, S. S., & Zhang, L. (2022). Single-cell transcriptomic and proteomic analysis of Parkinson’s disease Brains (p. 2022.02.14.480397). bioRxiv. 10.1101/2022.02.14.480397

